# Alpha and SSVEP power outperform Gamma power in capturing Attentional Modulation in Human EEG

**DOI:** 10.1101/2023.04.08.536134

**Authors:** Aritra Das, Nilanjana Nandi, Supratim Ray

## Abstract

The effect of visual attention has been extensively studied using various techniques such as macaque neurophysiology that yields spikes and local field potential (LFP), and human electro/magneto encephalogram (EEG/MEG). Attention typically suppresses power at low frequencies such as alpha band (8-12 Hz) and increases power in gamma band (>30 Hz) in brain signals. In addition, EEG studies often use flickering stimuli that produce a specific measure called steady-state visually evoked potential (SSVEP), whose power also increases with attention. However, effectiveness of these various neural measures in capturing attentional modulation is unknown since the stimuli and task paradigms vary widely across these studies. In a recent macaque neurophysiology study with flickering stimuli, we found that the effect of attention was more salient in the gamma band and beyond of the LFP, compared to alpha or SSVEP. To compare this with human EEG, we designed an orientation change detection task where we presented both static and counterphasing stimuli of matched difficulty levels to male (N=15) and female (N=11) subjects, allowing us to compare attentional modulation of various measures under similar conditions. We report two main results. First, attentional modulation was comparable for SSVEP and alpha. Second, non-foveal stimuli produced weak gamma despite various stimulus optimizations and showed a negligible effect of attention although full-screen gratings showed robust gamma activity. Our results are useful for brain-machine-interfacing studies where suitable features depending on recording modality are used for decoding attention, and also provide clues about the spatial scales of neural mechanisms underlying attention.

**Significance Statement:** Various neural measures such as alpha and gamma band power or SSVEP power capture signatures of visual attention. A systematic comparison of their effectiveness in capturing attentional modulation is important for understanding different neural computations underlying attention and for developing brain-machine-interfaces (BMIs) that can decode the focus of attention. Since stimulus configuration and task paradigms vary widely across visual attention studies, we recorded all the relevant neural signals during an attention task under similar stimulus and behavioral conditions in human EEG. In contrast to invasive recordings in which gamma outperforms other measures, our results show that in human EEG, attentional modulation of alpha and SSVEP are comparable, and non-foveal stimuli produce weak gamma which is not well modulated by attention.

## Introduction

Humans routinely use covert spatial attention in daily situations like searching for objects, playing sports, crossing the road, or driving. This mode of attention helps us overtly direct our attention to behaviorally relevant stimuli after selectively filtering out irrelevant information (Desimone and Duncan, 1995; Desimone, 1998). The neural correlates of covert spatial attention have been extensively studied using several techniques that record neural activity spanning different spatial scales, ranging from single/multi-unit spikes and local field potential (LFP) recorded using microelectrodes in multiple macaque visual areas, to diffuse population-level neural activity recorded using human electro/magnetoencephalogram (EEG/MEG). Attention-mediated modulation has been observed in evoked responses as well as in alpha (8-12 Hz) and gamma (>30 Hz) bands of the LFP activity in multiple visual areas in response to static/drifting stimuli (Fries et al., 2001, 2008; Chalk et al., 2010; Das and Ray, 2018) as well as in human EEG studies (alpha: Worden et al., 2000; Thut et al., 2006; gamma: Gruber et al., 1999; Müller et al., 2000) and MEG studies (Bauer et al., 2014; Magazzini and Singh, 2018). In addition, to address the poor signal-to-noise ratio in EEG/MEG recordings, human EEG studies often use flickering stimuli that produce a specific neural measure called steady-state visually evoked potentials (SSVEPs) to study the effect of covert spatial attention (Morgan et al., 1996; Kim et al., 2007; Itthipuripat et al., 2014b).

Since the stimuli and task paradigms vary widely across all these studies, previous studies have not attempted a thorough comparison of the effectiveness of these various neural measures in capturing attentional modulation. However, this comparison is important for two reasons. First, although earlier brain-machine interfacing (BMI) studies mainly focused on motor decoding (Bansal et al., 2012; Hwang and Andersen, 2013), recent studies have attempted to decode the focus of attention (De Sousa et al., 2021; Prakash et al., 2021; Chinchani et al., 2022) as well. For such applications, it is important to compare how attention modulates various neural measures. Second, different neural signals reflect important underlying neural mechanisms/computations operating over different spatial scales. For example, gamma oscillations are generated due to excitatory-inhibitory interactions (Buzsáki and Wang, 2012), especially involving soma-targeting parvalbumin positive interneurons (Bartos et al., 2007; Sohal et al., 2009), which remain coherent over a few millimeters (Jia et al., 2011; Murty et al., 2018). On the other hand, alpha oscillations could reflect thalamo-cortical gating mechanisms (Contreras et al., 1996). Recently, alpha has been linked with inter-areal feedback processing, while gamma is thought to reflect feedforward processing (van Kerkoerle et al., 2014; Michalareas et al., 2016). How attention modulates these measures therefore puts important constraints on the mechanisms underlying selective attention. For example, in a recent attention study in macaque V4 area, we found that the effect of attention was more salient in the gamma band and beyond of the LFP as compared to alpha or SSVEP (Prakash et al., 2021). We also compared the decodability of attentional state using spikes and LFP power at different frequency bands and found that gamma and high-gamma power recorded using a single electrode performed better than spikes; however, the decoding accuracy was comparable for high-frequency LFP (>30 Hz) and spikes when several electrodes were used. These results suggest that the spatial spread of mechanisms underlying attention could be comparable to the spatial spread of LFP (Prakash et al., 2022).

Here, we tested whether these results hold in human EEG, since it is a non-invasive recording technique routinely used in human BMI studies. Further, recording from a larger recording scale may also reveal important clues about the spatial scales of neural mechanisms of attention. To address this, we designed an orientation change detection task where we presented both static and counterphase flickering stimuli of matched difficulty levels, which allowed us to compare the attentional modulation of various measures under similar stimulus and behavioral conditions.

## Materials and Methods

### Human Subjects

A total of 28 subjects (16 Males and 12 Females, aged between 21-38 years, mean age of 26.10 years; standard deviation: 3.26 years) were recruited from the student community of the Indian Institute of Science for the EEG experiments. The Institute Human Ethics Committee of the Indian Institute of Science approved all the procedures before the start of the study. All the subjects reportedly had normal or corrected-to-normal vision, although none of the individuals were explicitly tested for visual acuity. The subjects voluntarily participated in the experiment after signing informed consent forms and were monetarily compensated after completing the experiments.

### EEG Recordings

Raw EEG signals were recorded from 64 active electrodes (actiCAP) using the BrainAmp DC EEG acquisition system (Brain Products GmbH). Electrodes were placed according to the international 10-10 system on a CAC-64 Brain Products EasyCap Layout. Electrolyte-gel (SuperVisc High Viscosity Electrolyte-Gel manufactured by EASYCAP GmbH) was applied to the scalp at each electrode contact site to lower the electrode impedances. Impedances of all electrodes were kept below 25 kΩ. Raw signals were filtered online between 0.016 Hz (first-order filter) and 250 Hz (fifth-order Butterworth filter), sampled at 1000 Hz, and digitized at 16-bit resolution (0.1 μV/bit). EEG signals were referenced to the FC_z_ electrode during data acquisition. We selected unipolar referencing for alpha, gamma, and SSVEP power estimation. We found that the bipolar referencing scheme, in which electrodes are re-referenced with respect to a neighboring electrode (see Murty et al., 2020, for details), gave slightly better gamma response than the unipolar referencing scheme for full-field gratings, similar to what has been reported by Murty and colleagues in earlier publications (Murty et al., 2018, 2020). However, for our dataset with non-foveal peripheral stimuli sets, the gamma responses were poor irrespective of the referencing scheme. Therefore, we used the unipolar referencing scheme for all our analyses.

### Experimental Design and Behavioral Tasks

Each subject was seated comfortably in a dark room in front of an LCD screen. Their heads were supported comfortably by a chin rest. The screen (BenQ XL2411) was gamma-corrected with a resolution of 1280 x 720 pixels and a refresh rate of 100 Hz. The screen was placed at a distance of 57 cm from the eye level of the subject. This arrangement allowed us to present full-screen gratings which subtended a width of ∼52° and height of ∼30° of visual field at an individual subject’s retina. All the visual stimuli were calibrated to the viewing distance of 57 cm. The stimuli were presented by custom software running on the MAC operating system, which controlled the task flow based on eye-tracking. The different types of behavioral tasks are summarized as follows:

#### Protocol 1: Gamma Tuning Protocol using Fixation Task

The goal of this protocol was to find the orientation and spatial frequency of a full-screen grating stimulus that produced the strongest gamma response. Every trial started with the onset of a white circular fixation spot (0.1°), presented at the center of the screen. The subjects were instructed to hold and maintain their gaze on that spot. After an initial blank period of 1000ms since the start of fixation, a series of stimuli (2-3) was presented for 800 ms each, with an interstimulus interval of 700 ms. These stimuli were full-screen sinusoidal luminance gratings presented in full (100% luminance w.r.t to gray background) contrast. During stimulus presentation, the gratings were either static or flickering at 16 Hz counterphase. The spatial frequency (SF) of the gratings could take either of the two values: 2 or 4 cycles per degree (cpd), whereas the orientation of the gratings could be any of the four values: 0°, 45°, 90°, and 135°. We chose these stimulus parameters as they were shown to produce robust gamma responses previously (Murty et al., 2018, 2020). Stimuli were presented pseudo-randomly to avoid any possible adaptation effects. Subjects performed this task for 15-20 minutes with each of the eight combinations (2 SFs x 4 Orientations) presented pseudo-randomly 20-30 times (Refer to Figure 1A). The subject’s fixation was monitored online by EyeLink 1000 (SR Research Ltd, sampled at 500 Hz) during the entire trial period. Characteristic auditory feedback was provided to the subject for trial onset, fixation onset, trial abortion due to fixation break, or correctly completing a trial by maintaining fixation. Twenty-eight subjects (16 Males and 12 Females) performed this task at the beginning of a recording session. We determined the spatial frequency and orientation of grating stimuli which generated the maximum gamma band power response (25-70 Hz; see details of power estimation below). We used this spatial frequency and orientation for the remaining protocols.

**Figure 1:**
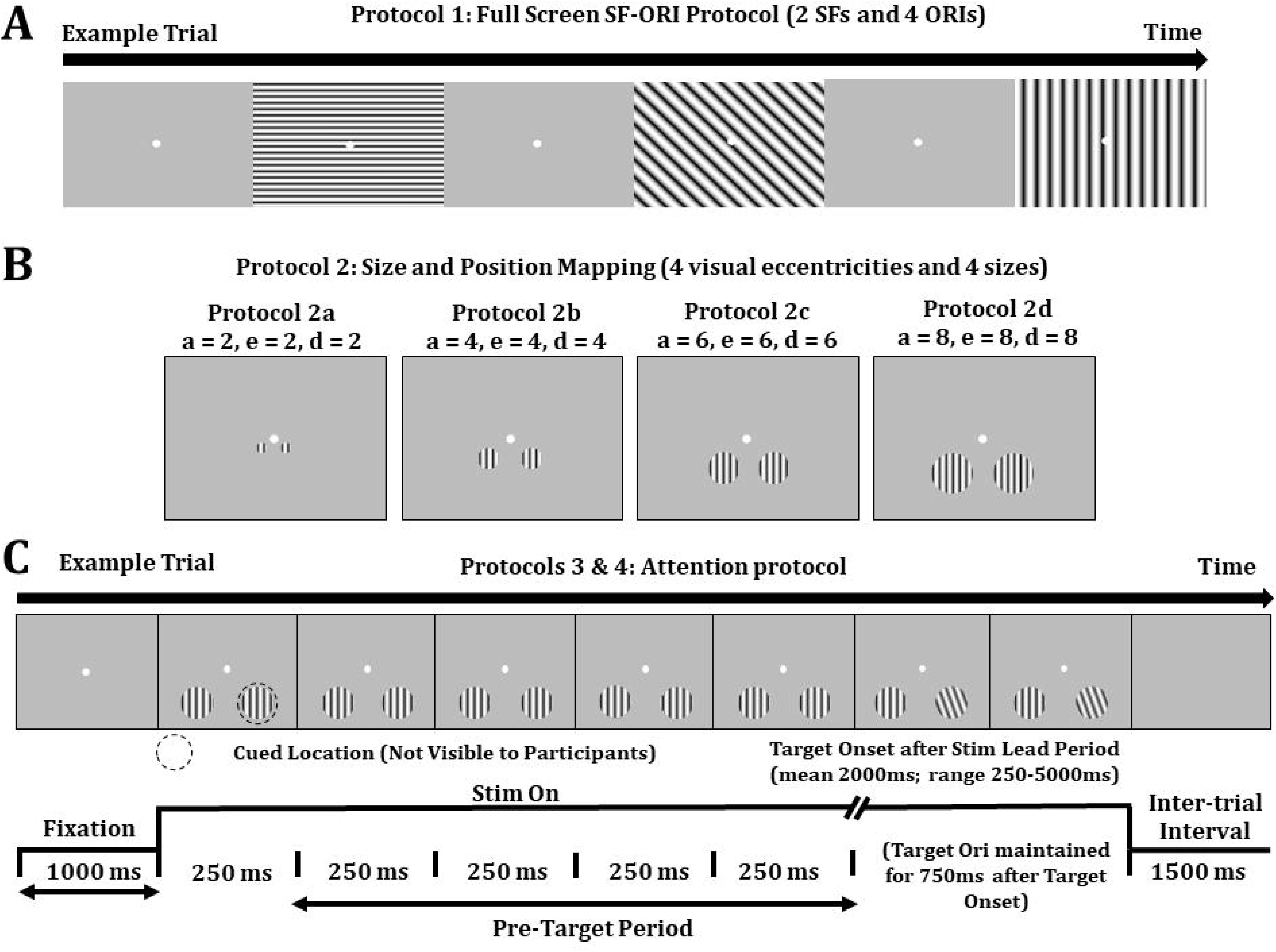
Task Paradigms for human EEG experiments. (A) Gamma Tuning Protocol: While the subjects maintained passive fixation, full-screen grating stimuli were presented at full contrast for 800ms with an inter-stimulus interval of 700 ms, with 2-3 stimuli shown in each trial. During a stimulus presentation, stimuli could either be static or counterphase flickering at 16 Hz. The values of spatial frequency (SF) were either 2 or 4 cpd, and orientation values could take four possible values (0°, 45°, 90°, and 135°). (B) Stimulus Size and Eccentricity Mapping Protocol: While the subjects maintained passive fixation, two equidistant non-foveal achromatic gratings were presented peripherally in the lower visual field while the diameter and eccentricity of the gratings were varied systematically in separate experiments (protocol 2a-2d). The azimuth and elevation of the center of the gratings and the diameter were varied from 2°-8° in steps of 2° in protocols 2a-2d. During each trial in any of the protocols, the pair of stimuli were presented at full contrast for 800 ms with an inter-stimulus interval of 700 ms, with 2-3 stimuli shown in each trial. During stimulus presentation, either both the gratings were static or the left grating counterphase flickering at 12 Hz and the right grating flickering at 16 Hz. The spatial frequency and orientation values for an individual subject were determined based on the SF-Ori combination that produced the maximum gamma response (25-70 Hz) for static gratings in the gamma tuning protocol described in A. (C) Attention Task: While the subjects maintained fixation, two achromatic gratings were presented at equidistant spatial locations from the fixation spot in the lower visual field. The subjects were cued to covertly attend to one of the two locations in different blocks of trials (indicated by a black dotted circle for clarity; not visible to the subjects). Grating parameters (orientation and spatial frequency) were determined for each subject based on A. The diameter, azimuth, and elevation for all the subjects were set to 4° (Protocol 2b), based on results of B on a group of 11 subjects before performing any attention experiments. During each trial, either both the stimuli were static, or both were counterphase flickering, one at 12 Hz and the other at 16 Hz. The contrasts of both the attended and ignored locations were at 100% during each trial. There was an initial time period of 1000ms at the beginning of each trial when the subject maintained their fixation, followed by an unstipulated time before the orientation of the stimulus at the cued location changed (target). The stimuli were considered to be 250 ms long, which contained 3 and 4 full cycles of the 12 and 16 Hz stimuli, presented without any inter-stimulus interval so that they appeared to be flickering continuously. The target appeared at an unstipulated time drawn from an exponential distribution (mean 2000 ms, range 250-5000 ms) following the initial stimulus lead period of 1000 ms. The subjects had to make a saccade to the target location within 800 ms of the target onset for a correct trial (hit). If there was no change during a trial (catch trial), the subjects had to maintain their fixation. In protocol 3, each subject performed this attention task for 2-3 blocks on each side where the orientation change of the cued gratings (target) could occur randomly from any of 5 preset values. The preset orientation change values were different for each of the temporal frequencies (static, 12, and 16 Hz). The psychometric curves for each static and flickering gratings were fitted separately with Weibull functions after pooling the behavioral accuracy values from both sides. The orientation change values for 50% accuracy for each of the static and flickering gratings were determined from the fits. This single target orientation (separate for static, 12 and 16 Hz) was used for protocol 4 in which the subjects performed 5-6 blocks on each side.

#### Protocol 2: Stimulus Size and Eccentricity Mapping using Fixation Task

Since covert spatial attention tasks typically use two stimuli presented equidistant from a central fixation spot, we had to find the optimal stimulus eccentricity and size to elicit robust neural responses in EEG. The size and eccentricity of non-foveal stimuli can profoundly affect behavior and neural responses. If the stimuli are too big, it would be easier to detect orientation changes in an attention protocol and subjects may only attend to the portion of the stimulus; therefore, the attentional modulation of neural signals may be lower. On the other hand, if the stimuli are small, they may not elicit a robust neural response in EEG. Similarly, the covert attention task would become easier if the stimuli are presented near the fixation spot (lower eccentricity). But if stimuli are presented too far from the fixation spot, they may not produce a robust response, or, in our attention task design (described below), it could be difficult for subjects to make eye movements toward the target within a specified time.

We therefore tested several possible combinations of stimulus size and eccentricity to find a combination that produced robust neural signals in EEG. Every trial started with the onset of a white fixation spot (0.1°). We presented two achromatic grating stimuli located equidistant from the central fixation spot in the lower visual hemifield with the azimuth and elevation of the center of the two stimuli at (−;X°, −Y°) and (X°, −Y°), with the diameter being D° of visual angle. In four consecutive experiments, we varied the values of X, Y, and D set at 2°, 4°, 6°, or 8° (refer to Figure 1B). The eccentricities of the four non-foveal stimuli sets, i.e., the distance between the center of the stimulus and the center of vision (point of fixation or fovea), were 2.8, 5.6, 8.5, and 11.3°, respectively. After an initial blank period of 1000 ms after the start of fixation, a series of stimuli (2-3) was presented for 800 ms each, with an interstimulus interval of 700 ms. These stimuli were full-screen sinusoidal luminance gratings presented in full (100% luminance w.r.t to gray background) contrast. During each stimulus presentation, either both the gratings were static, or the left and right gratings were counterphase flickering at 12 Hz and 16 Hz, respectively, to mimic the experimental paradigm for the attentional task later. We presented both the stimuli in the lower visual hemifield because the occipital electrodes placed on the scalp are directly below the region of the primary visual cortex above the calcarine fissure in the occipital cortex of the human brain, and they are known to be retinotopically mapped to the visual cortex from the lower visual field. The subjects performed each of the four protocols for 5-10 minutes with 25-30 repeats each for static and flickering stimuli presentations.

The first two protocols were run on 11 subjects (7 Males and 4 Females) before any attention task (Protocols 3 and 4; see below) were done. We found that X, Y, and D of 4° gave the best response for SSVEPs, while the responses for alpha and gamma did not depend on the eccentricity (Fig 3). Therefore, we fixed this stimulus configuration for all subjects. Attention task on these 11 subjects was done on a different day. For the remaining subjects, Protocol 1 was followed by Protocols 3 and 4 and all three protocols were done on the same day.

#### Protocols 3 and 4: Attention Task

We finally used the spatial frequency, orientation, size, and eccentricity optimized for the attention task, where we presented both static and flickering stimuli to measure all the relevant neural measures. Subjects were required to maintain their gaze within 2° of a small central dot (diameter of 0.10°) during the task while covertly attending to one of the two achromatic grating stimuli presented in the lower hemifield equidistant from the fixation spot. The subjects were cued to one of the two stimulus locations (left or right) in different blocks of trials by presenting an instruction trial at the start of the block, in which there was only a single stimulus either on the left or right of the fixation spot, indicating the location to be attended in subsequent trials of that block. During each trial, either both the stimuli were static, or both were counterphase flickering, one at 12 Hz and the other at 16 Hz. We used 12 Hz and 16 Hz frequencies since these had 3 and 4 complete cycles during a 250ms interval. In our protocol, one “stimulus” was considered to be of 250 ms, and a sequence of such stimuli was presented without any interstimulus interval, and therefore appeared as a continuous stimulus flickering at 12 or 16 Hz. The contrasts of both the attended and ignored locations were at 100% in each trial. There was an initial time period of 1000ms at the beginning of each trial when the subject maintained their fixation, followed by the stimulus onset. Following stimulus onset, the orientation of the stimulus at the cued location changed (target). The target appeared at an unstipulated time drawn from an exponential distribution (mean 2000 ms, range 250-5000 ms) following the initial stimulus lead period of 1000 ms. The initial stimulus lead period was used to avoid transient activity generated by the event-related potential during the 250 ms after the first stimulus onset. An exponential distribution was used for target appearance to minimize the expectation of target appearance and to keep the attentional state uniform during a trial since the hazard function is flat for an exponentially distributed target onset time. Subjects were required to make a saccadic eye movement towards the target within 800 ms of the target onset. Once the target appeared, it remained at the changed orientation for 800 ms. To account for saccade latency and minimize guessing, saccades beginning at least 100 ms after the target appearance were considered correct trials. We also introduced catch trials in which a trial was terminated at 6250 ms if the target had not appeared till then (15-20% of all trials). Characteristic auditory feedback was provided to the subjects for either correctly detecting targets in a trial (Hit) or for maintaining fixation throughout catch trials. A different auditory sound was provided as feedback for failing to detect the target (Miss) or fixation breaks. For the non-catch trials, targets appeared at the cue location (cue validity 100%; no invalid trials). This task design was used to rule out any distributed attention strategy the subjects may use, which can influence the neural responses.

Note that our task design made the detection of a target for the static case much easier than the flickering case. For counterphase flickering stimuli, the contrast varied sinusoidally to complete 3 or 4 cycles within a 250 ms “stimulus period”, so the target stimulus (with a different orientation) appeared initially at 0% contrast at the beginning of the target stimulus onset (in other words, the orientation change occurred when the stimulus contrast was 0%). This was not the case for static stimuli, making them much easier to detect. Therefore, to match the difficulty levels of the static and flickering stimuli, we performed 2-3 blocks of attention task with five different orientation changes for each of the static, 12 Hz, and 16 Hz stimuli for each of the two attended locations (left and right). Each block comprised of 10 trial repeats for both hit and miss trials combined. We plotted the behavioral accuracy for the two attended locations for all five orientation changes. We fitted the combined behavioral performance for both left and right locations with a Weibull function for psychometric analysis. We determined the fitted orientation change value for each temporal frequency (TF; static, 12, and 16 Hz), for which the behavioral accuracy was at 50%.

We then set the single orientation change value obtained from the above manual behavioral staircase procedure for each of the TFs for our main attention task. The subjects again performed the main attention task for six blocks for each of the two attended locations where either both static or both flickering stimuli were presented in each trial. Each block comprised of 10 trial repeats for both hit and miss trials combined. We had at least 100-120 trial repeats for each TF with hits and misses conditions combined for each subject. Two subjects (one male and one female) were unable to perform the attention task. Therefore, the attention dataset had 15 male and 11 female subjects.

### Data Analysis

All data were analyzed using custom codes written in MATLAB (The Mathworks, RRID: SCR_001622). Individual data analysis methods are briefly summarized below.

#### EEG Artifact Rejection and Electrode Selection

We followed a standard set of procedures for electrode selection and artifact rejection in human EEG experiments reported previously in detail (Murty et al., 2020; Murty and Ray, 2022). We briefly describe the procedure here with a minor modification. To find bad stimulus repeats for each electrode, we applied a thresholding procedure on raw waveforms (high pass filtered the signal at 1.6 Hz to eliminate slow drifts) and multi-taper PSD in the analysis period (Fixation Protocols: −500 ms to 750 ms of stimulus onset; Attention Protocol: combined periods of −1000 ms to 1250 of the first stimulus onset in a trial and −1000 to 0 ms of target onset) using the Chronux toolbox in MATLAB (Bokil et al., 2010). We labeled a particular stimulus repeat as bad if the RMS value of the time-series signal was beyond a pre-specified threshold (min: 1.5 µV, max: 35 µV) or PSD of that stimulus repeat deviated by six times the standard deviation from the mean at any frequency bin between 0 and 200 Hz for a given electrode. The only difference in EEG artifact rejection in this study compared to previous studies is the usage of RMS threshold for the time-series signal instead of the standard deviation-based threshold around the mean of the time-series signal reported in our previous EEG studies (Murty et al., 2020; Murty and Ray, 2022). If an electrode had more than 30% bad repeats, we discarded that electrode. We then created a set of common bad stimulus repeats across the remaining electrodes if a particular repeat was labeled as bad in more than 10% of those remaining electrodes. Finally, any repeat that was labeled bad in any of the ten electrodes (P3, P1, P2, P4, PO3, POz, PO4, O1, Oz, and O2) was added to the pool of common bad repeats, thus finalizing the list of common bad stimulus repeats. In addition, we removed electrodes with baseline PSD slopes less than zero in the gamma range of 56-84 Hz (for details, please refer Murty et al., 2020). Using this set of stringent criteria, we rejected ∼20% of data for fixation protocols, similar to previous reports by Murty and colleagues. However, this increased to ∼30% when we considered attention datasets where we checked bad trials for the combined periods of −1000 ms to 1250 of the first stimulus onset in a trial and −1000 to 0 ms of target onset, since two epochs were considered instead of one. For all the experiments, we analyzed occipital eight electrodes (P1, P3, PO3, and O1 on the left hemisphere and P2, P4, PO4, and O2 on the right hemisphere). Previous EEG studies from our group have also used the similar set of occipital electrodes to characterize alpha and gamma oscillations (Murty et al., 2018, 2020) and SSVEP response (Murty et al., 2020; Liza and Ray, 2022).

#### EEG data analysis

Power spectral densities (PSDs) for different stimulus or attention conditions were computed using the multi-taper method with one Slepian taper using the Chronux toolbox (Bokil et al., 2010; https://chronux.org/, RRID: SCR_005547).

For Figures 2B and 2D, the analysis period was selected between 250 ms to 750 ms after stimulus onset to avoid stimulus onset-related transients and compared against a ‘baseline period’ between −500 and 0 ms of stimulus onset (a gray screen). This yielded a frequency resolution of 2 Hz for the PSDs. Absolute PSDs were expressed in logarithmic units (base 10), and the changes in PSDs were plotted with respect to the baseline response:

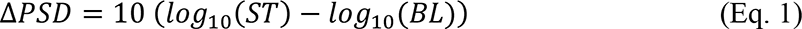

**Figure 2:**
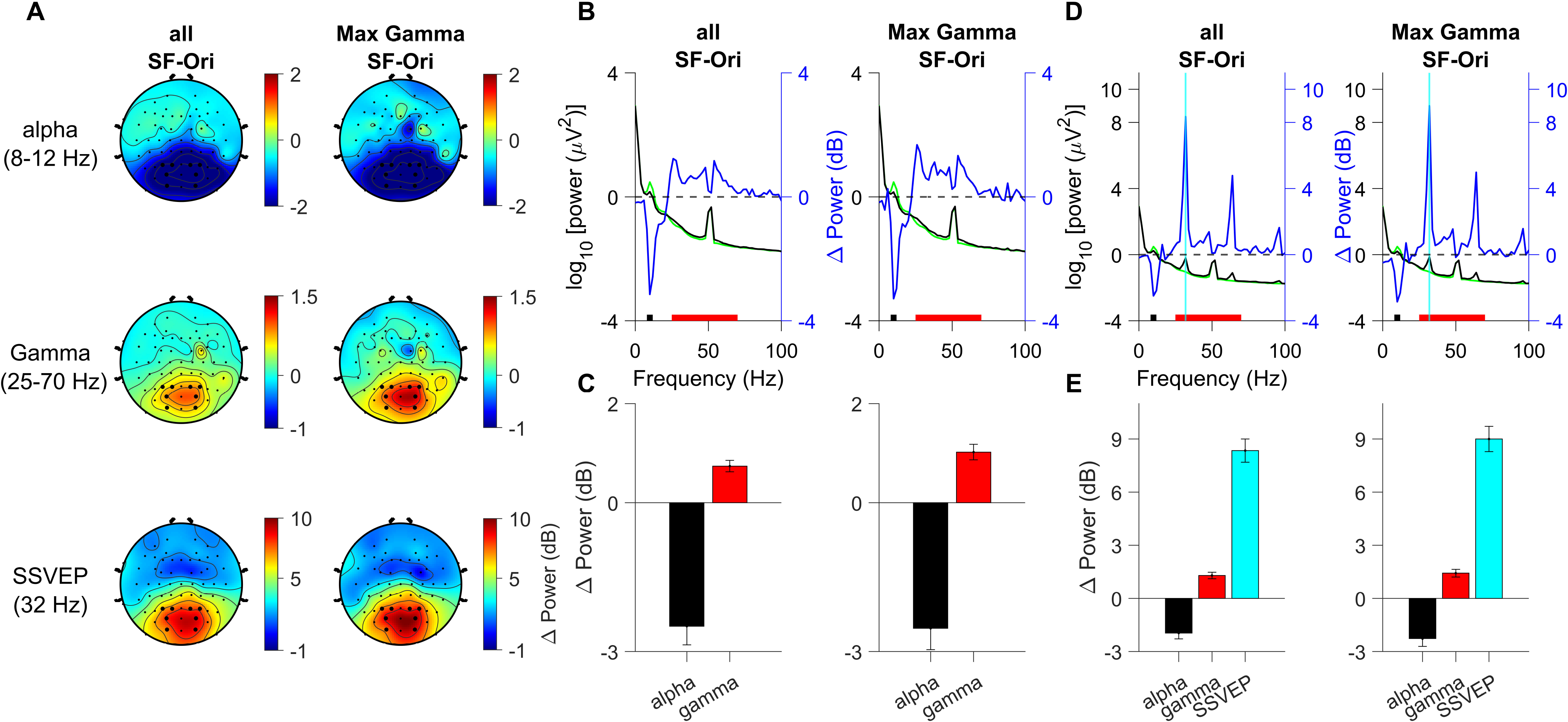
alpha, gamma, and SSVEP response in subjects for full-field grating stimuli. (A) Scalp maps showing 64 electrodes (represented as dots). Scalp maps show the change in alpha power (top row) in the stimulus analysis period (250 to 750 ms) w.r.t baseline (−;500 to 0 ms; zero denotes stimulus onset), gamma (middle) for static gratings, and SSVEP power (bottom) computed by the trial-by-trial power estimation method averaged across 28 subjects. The power changes are separately shown for all SF-Ori combinations (left column) and the SF-Ori combination that produced maximum gamma response (right column). Eight analysis electrodes used in Panels B-E are marked in bold dots. (B) Left ordinate (black) shows raw power spectral densities (PSDs) vs. frequency for stimulus (250 to 750 ms, black trace), baseline (−;500 to 0 ms, green trace) analysis periods averaged across all 28 subjects for the eight electrodes highlighted in panel A for all SF-Ori combinations (left), and max SF-Ori combination (right) in response to static gratings; right ordinate (blue) shows the same for change in PSD (in dB, solid blue trace). Solid black and red bars on the X-axis indicate alpha (8-12 Hz) and gamma bands (25-70 Hz), respectively. (C) Change in alpha (black bar), gamma (red) power in the stimulus analysis period (250 to 750 ms, black trace) w.r.t baseline (−;500 to 0 ms, green trace) analysis period averaged across all 28 subjects for the eight electrodes highlighted in panel A for all SF-Ori combinations (left), and max SF-Ori combination (right)) in response to static gratings; The respective powers were calculated in the frequency ranges highlighted in panel B. Error bars indicate the standard error of the mean across subjects (N=28). (D) Same layout as in B in response to 16 Hz counterphase flickering gratings. The SSVEP response at 32 Hz (twice that of the flickering frequency) is highlighted by a solid cyan vertical line. (E) Same layout as in C in response to 16 Hz counterphase flickering gratings with the addition of change in SSVEP power (cyan bar).

where ST is stimulus power spectra (across frequency f) averaged across repeats for all eight SF-Ori combinations (two SFs and four orientations) or stimulus combination that produced the maximum gamma response and all analyzable electrodes. BL denotes the baseline power spectra averaged across repeats for all stimulus combinations and analyzable electrodes.

We calculated the change in alpha and gamma power (Figures 2 and 3) as follows:

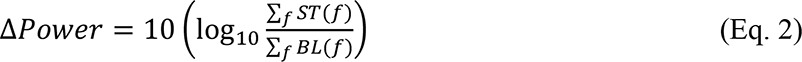

where ST (all/max gamma stimulus combination) and BL (all) are stimulus and baseline power spectra (across frequency f) averaged across repeats and analyzable electrodes. For alpha, f ∈ [8 12] Hz, and for gamma, f ∈ [25 70] Hz. For changes in SSVEP power, we calculated the power at the 2^nd^ harmonic of the counterphasing frequency. The fundamental frequency response (the first harmonic of the flickering frequency) in the PSD, generated due to activity of cortical simple cells, mostly averages out due to the random spatial distribution of the ON and OFF sub-regions of their receptive fields with respect to the symmetric luminance flickers of counterphase flickering gratings (Campbell and Maffei, 1970; De Valois et al., 1982; Di Russo et al., 2001; Hou et al., 2003). The SSVEP response at second harmonic (f_2_ = 2f_1_) arises due to activity of the frequency doubling cortical complex cells (De Valois et al., 1982; Di Russo et al., 2001; Kim et al., 2007) whose responses are largely spatial-phase invariant.

For attention results in figures 5 (static gratings), and 7 (flickering gratings), the analysis period was selected for the pre-target period between −1000 ms to 0 ms of target onset and compared against a ‘baseline period’ between −1000 and 0 ms of first stimulus onset (a gray screen). This yielded a 1 Hz frequency resolution in PSDs. The change in PSDs (violet traces in figures 5C, and 7B) between attended and ignored conditions were computed as follows:

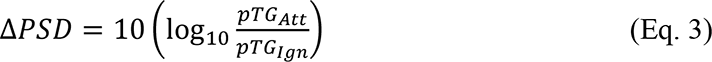

Where pTG_att_ and pTG_Ign_ are pre-target power spectra for attended and ignored conditions, respectively (across frequency f), averaged across trials (static and flickering trials separately), attention conditions, and analyzable electrodes.

We calculated the change in power in attended and ignored conditions w.r.t. to baseline period using the following equation:

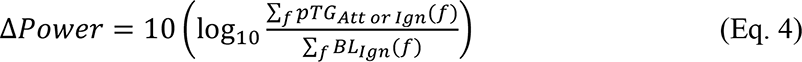

Where pTG and BL are pre-target, and baseline power spectra (across frequency f) were averaged across either static grating conditions (Figure 5A-B ‘Attend Left’ and ‘Ignore Left’ scalp maps) or flickering grating conditions (Figure 7A ‘Attend Left’ and ‘Ignore Left’ scalp maps) for alpha, f ∈ [8 12] Hz, and for gamma, f ∈ [25 70] Hz ranges. For flickering gratings, the power at SSVEP harmonics were excluded during computation of gamma band power. For changes in SSVEP power, we calculated the power at the frequency twice that of the counterphasing frequency (i.e., 24 Hz or 32 Hz depending on the attended or ignored grating flickering at 12Hz or 16 Hz, respectively). We use equation 4 to calculate power change in bar plots, with pTG and BL being the pre-target and baseline power spectra (across frequency f), which were averaged across all trials (static and flickering trials separately), attention conditions, and analysis electrodes. For delta PSDs (Figure 5C and Figure 7B) and bar plots (Figure 5D and 7C), we computed the change in PSD/power between attend-left/ignore-left conditions for the four right-hemispheric occipital electrodes (P2, P4, PO4, and O2) and for attend-right/ignore-right conditions for the four left-hemispheric occipital electrodes (P1, P3, PO3, and O1). Please note that the same baseline power is subtracted from both attention conditions to rule out the possibility that the results are due to changes in baseline power.

All the absolute PSDs, relative PSDs, and change in power values computed for an individual subject (equations 1-4) were subsequently averaged across subjects, whereas, for scalp maps of 64 unipolar electrodes, each analyzable electrode data was averaged across analyzable repeats/trials, attention conditions and subjects (Figures 2, 3, 5-7). Please note that we also removed the three frequency bins centered around the line noise peak (52 Hz; Figures 2 and 3) and five frequency bins centered around the line noise peak (51 Hz; Figures 5 and 7) while calculating gamma power to remove the line noise artifact.

#### Scalp Maps (Figure 2, 3, 5-7)

Unlike the scalp maps for alpha, gamma, and SSVEP in the gamma tuning protocol (Figure 2) and alpha and gamma in the size-eccentricity protocol (Figure 3), we consolidated the SSVEP responses for our size-eccentricity protocol (Figure 3A; bottom two rows) and attention protocol (Figure 7A; bottom two rows) for a simple and effective comparison across different neural measures by presenting SSVEP responses on the right hemispheric occipital electrodes in the consolidated scalp maps. To achieve this, we mirror-flipped the scalp map for the 32 Hz SSVEP response for 16 Hz counterphasing grating presented on the lower right visual hemifield in our size-eccentricity mapping protocols with respect to the midline set of electrodes (z-line of ActiCap-64 Cap) and averaged with the 24 Hz SSVEP response for the 12 Hz counterphasing grating presented on the lower left visual hemifield. Similarly, for the attention protocol, the SSVEP responses for all four attention conditions (two attended locations and two TFs; figure 6A-D) were averaged after mirror-flipping the scalp maps for two attend-right (12 Hz/ 16 Hz) conditions. For the alpha and gamma scalp maps for flickering gratings, the scalp maps were averaged across only attend-left 12 Hz and attend-left 16 Hz conditions.

**Figure 3:**
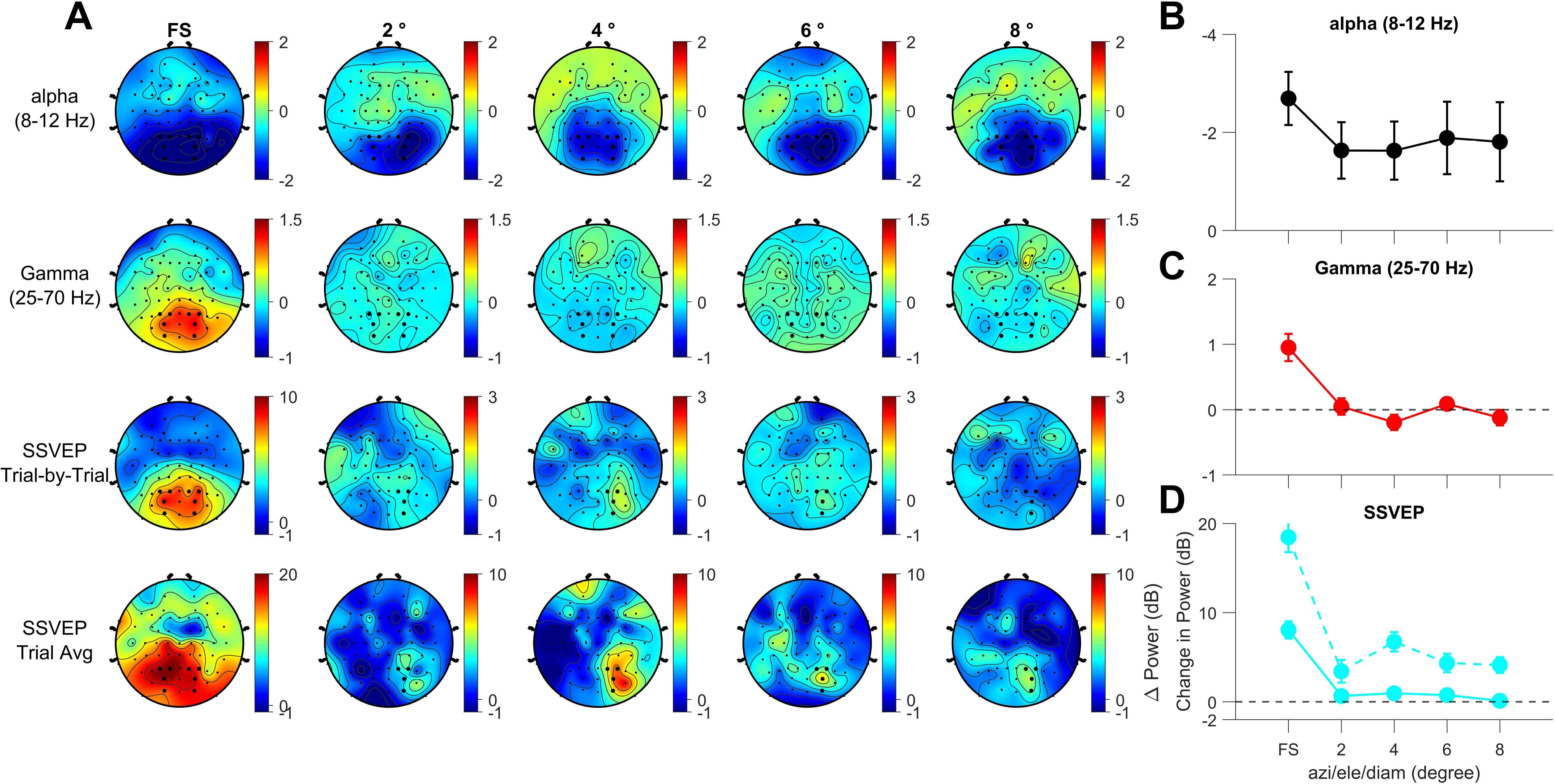
alpha, gamma, and SSVEP response in subjects in response to non-foveal stimuli pair. (A) Scalp maps showing 64 electrodes (represented as dots). Color maps show the change in power in the stimulus analysis period (250 to 750 ms) w.r.t baseline (−;500 to 0 ms) (top row) for alpha, gamma, SSVEP power using trial-by-trial SSVEP power estimate, and by the trial-averaged SSVEP power estimate from top to bottom rows averaged across 11 subjects. The columns indicate the diameter and stimulus center locations of the gratings used in separate experiments in protocol 1 and protocols 2a-2d mentioned in Figure 1. Eight electrodes used in Panels B-D are marked as bold dots. Note that while all 8 electrodes are used for alpha and gamma calculation, the calculation is done differently for SSVEPs (protocols 2a-2d). Here, since the two stimuli flicker at two different frequencies (12 and 16 Hz), the SSVEP response for a particular frequency is measured only for the contralateral 4 electrodes. The SSVEP plots shown here are obtained by first mirror-flipping the scalp map for the stimulus on the right side and then averaging with the scalp map for the left stimulus (as if the stimulus is always on the left side), which is why the SSVEP responses are lateralized on the right side. Consequently, only the 4 contralateral electrodes are highlighted in these plots. Note, however, that these 4 electrodes represent averaged data from all 8 electrodes, since data from the 4 left electrodes are mirror-flipped to the right side. (B-D) Change in power in alpha (B), gamma (C), and SSVEP (D) for stimulus (250 to 750 ms, black trace), baseline (−;500 to 0 ms, green trace) analysis periods averaged across all 11 subjects for the electrodes highlighted in panel A. Note that, the ordinate of panel B is flipped upside down to highlight the magnitude of alpha suppression. Panel D shows trial-by-trial SSVEP power estimates (solid cyan lines) and trial-averaged power estimates (dotted cyan lines).

#### SSVEP power calculation

We computed SSVEP power in two different ways, which are as follows:

**Method 1:** We computed the power for each trial and subsequently computed the mean of the power across all trials. We termed this the **trial-by-trial SSVEP power estimate** (Itthipuripat et al., 2014b; Murty et al., 2020; Salelkar and Ray, 2020).

**Method 2:** Averaged the EEG signal across all trials and then computed the power from the averaged signal. We called this the **trial-averaged SSVEP power** (Morgan et al., 1996; Liza and Ray, 2022).

#### Statistical Analysis

Statistical tests were performed for each of the three neural measures to check whether attention significantly modulated these measures or whether the magnitude of these modulations differed between different pairs of neural measures. We used pairwise t-tests to compare the magnitude of attentional modulation on different neural measures (Figures 5D and 7C). Since gamma and SSVEP showed an increase in attentional modulation whereas alpha showed suppression with attention, for statistical comparisons of magnitude of attentional modulations between alpha and other neural measures, we changed the signs of alpha power changes and then performed pairwise t-tests. We also performed t-tests for each neural measure to check whether they were significantly modulated by attention.

## Results

### Full-field stimuli produce robust alpha, gamma, and SSVEP responses in human EEG

We first analyzed the three neural measures (alpha, gamma, and SSVEP power) by recording EEG signals for full-field stimuli presented to subjects passively fixating at the central fixation dot. We recorded full-field stimulus responses for both static as well as flickering stimuli for 28 subjects and analyzed the results for eight electrodes (Electrodes P3, P1, PO3, and O1 on the left hemisphere and P2, P4, PO4, and O2 on the right hemisphere) referenced to the FCz electrode. We computed the raw power separately between 250-750 ms after stimulus onset and a baseline period of −500 ms to 0 ms of stimulus onset for all the electrodes of an individual subject. We averaged the power across the occipital electrodes and computed the power changes in decibels for an individual subject, followed by averaging the power changes across all the subjects.

Figure 2A left column shows the scalp maps separately for alpha, gamma, and SSVEP responses for all SF-Ori combinations for all subjects. All sixty-four electrodes are shown as dots in scalp maps, with the analysis electrodes highlighted as bold dots. The full-field stimuli produced strong alpha suppression (∼ −2 dB) and robust gamma power (∼1 dB) for static stimuli in the parietal, parieto-occipital, and occipital electrodes. We also found strong SSVEP power (trial-by-trial estimate; see Methods for details) at 32 Hz (∼8 dB), the second harmonic of the 16 Hz counterphase flickering stimuli. The scalp maps in Figure 2A right column show the alpha, gamma, and SSVEP responses for the spatial frequency and orientation that produced the maximum gamma response for the eight analysis electrodes. As expected, we observed an increase in gamma power in this stimulus combination. Alpha and SSVEP responses expectedly remained almost similar for the two conditions (left and right panels in A) since the stimulus combination was chosen to maximize the gamma response.

We computed the PSDs of the eight analysis electrodes in Figure 2B for static stimuli, which show suppression in the alpha band for the stimulus period (black trace) with respect to the baseline period (green trace). The blue trace indicates the change in PSDs. The PSDs further show an increase in gamma band power for all SF-Ori and max Gamma SF-Ori combinations for static gratings (Figure 2C) and an increase in SSVEP power for flickering gratings (Figure 2D). We quantified alpha and gamma power for static stimuli and SSVEP for flickering stimuli in Figures 2C and 2E, respectively. Even though we observed an increase in gamma power for flickering stimuli, the PSDs show that the counterphase stimuli produced strong SSVEP signals at the second harmonic and all subsequent even harmonics, thus interfering with gamma response. Therefore, for the gamma power calculation in Panel E, we removed the power at SSVEP harmonics. Taken together, the magnitude of SSVEP power was much higher (∼8-9 dB) compared to the magnitude of alpha suppression (∼2 dB) and gamma (∼1-1.5 dB). These results were consistent with previous EEG studies (Murty et al., 2018, 2020).

### Comparison of alpha, gamma, and SSVEP responses in human EEG for varying size and eccentricity of non-foveal stimuli

Next, we used the particular spatial frequency and orientation that produced maximum gamma response for the size and eccentricity mapping protocols in 11 subjects. We systematically varied both the azimuth and elevation of the stimulus center as well as the diameter of the stimulus from 2° to 8° in steps of 2 degrees of visual angles in four consecutive experiments. We show the scalp maps for the alpha, gamma, and SSVEP power for these different experiments in Figure 3A, along with the scalp maps for full-screen stimuli in the leftmost column for comparison. We found robust alpha suppression for all sets of non-foveal stimuli, although it was reduced compared to full-screen gratings in the occipital electrodes. The alpha suppression decreased in the non-foveal gratings by about 0.4-0.6 dB (Figure 3B).

For gamma power, non-foveal stimuli produced a negligible response despite showing a robust gamma response of ∼1 dB in full-field stimuli (Figure 3A gamma scalp maps and Figure 3C). In an earlier study on humans with size tuning protocols, Murty and colleagues had varied the diameter of foveal gratings from 1°-32° in steps of 2° in addition to full-screen gratings (>32°). They found the magnitude of gamma response progressively increased with increasing stimulus size, and they also found two distinct bands, slow (20-40 Hz) and fast (40-70 Hz), from a diameter of 4° onwards (Murty et al., 2018). However, we did not observe gamma responses in either band when we separately analyzed both slow and fast gamma bands for our non-foveal stimuli (data not shown).

Next, we computed the SSVEP power for each electrode of an individual subject in two ways: 1) trial-by-trial SSVEP power computed for each stimulus repeat and subsequently averaged across repeats, and 2) trial-averaged SSVEP power computed by averaging the signal across stimulus repeats and then computing the power (see Methods). The trial-averaged SSVEP is often used in EEG studies to estimate SSVEP power better because averaging the trials before power calculation enhances the signal-to-noise ratio at the SSVEP frequency (second harmonic and all subsequent even harmonics). For the trial-by-trial SSVEP power estimate, the best SSVEP response was ∼1 dB in the analysis electrodes for stimuli presented for location and diameter of 4° compared to other non-foveal stimuli sets, although it was much smaller than the full-field response (Figure 3D). The SSVEP response, however, increased to about ∼6 dB for diameter and location of 4° when computed by the trial-averaged method. Note that we combined the SSVEP responses at 24 Hz from the right-side analyzable electrodes for the 12 Hz flickering stimuli on the left and SSVEP responses at 32 Hz from the left-side analyzable electrodes for the 16 Hz flickering stimuli on the right. For scalp maps of SSVEP response, we mirror-flipped the scalp maps for 32 Hz along the midline of our cap layout to show on the right hemisphere, which is why the SSVEP response appears lateralized to the right side. Based on these results from 11 subjects, we selected the grating location and diameter of 4° for attention tasks because it showed the maximum SSVEP response and robust alpha suppression.

### Behavioral Results for Attention Task

The difficulty levels for static and flickering stimuli in the attention task were matched by fitting psychometric curves for behavioral performances for a 2-3 block pilot attention experiment for each of the attended sides (left or right) with five possible values of orientation change. A single orientation change value for each of the temporal frequencies (static, 12 Hz, and 16 Hz) of the cued location was obtained from the fitted values for which the behavioral performance was at 50%. We used these orientation changes for static and flickering stimuli in the main attention task (see methods section). Subjects performed at least five to six blocks on each attended side (left or right). In Figure 4, we present the behavioral performances of all 26 subjects who performed the main attention task. Behavioral accuracies were similar in all attended locations and temporal frequencies, except for the att-left 16 Hz condition (Figure 4A). The behavioral accuracy for att-Left 16 Hz, when compared to other attention conditions, was significant (all the p values < 0.05) whereas the behavioral accuracy was not significantly different between the other attention conditions (all p values > 0.18). The orientation thresholds (Figure 4C) were set to yield a behavioral performance of 50%, but the actual performance was slightly better (∼70% on average), potentially due to practice and familiarization with the task paradigm. Response times for static gratings (both left and right attended locations) were lower than for flickering gratings (Figure 4B). The pairwise t-tests between the response time between any static and flickering stimuli were significant (all the p values < 1.12 × 10^-5^). Orientation change values (target) were considerably smaller (∼0.6-1°) for static as compared to flickering stimuli (∼10°), as described in the Methods section.

**Figure 4:**
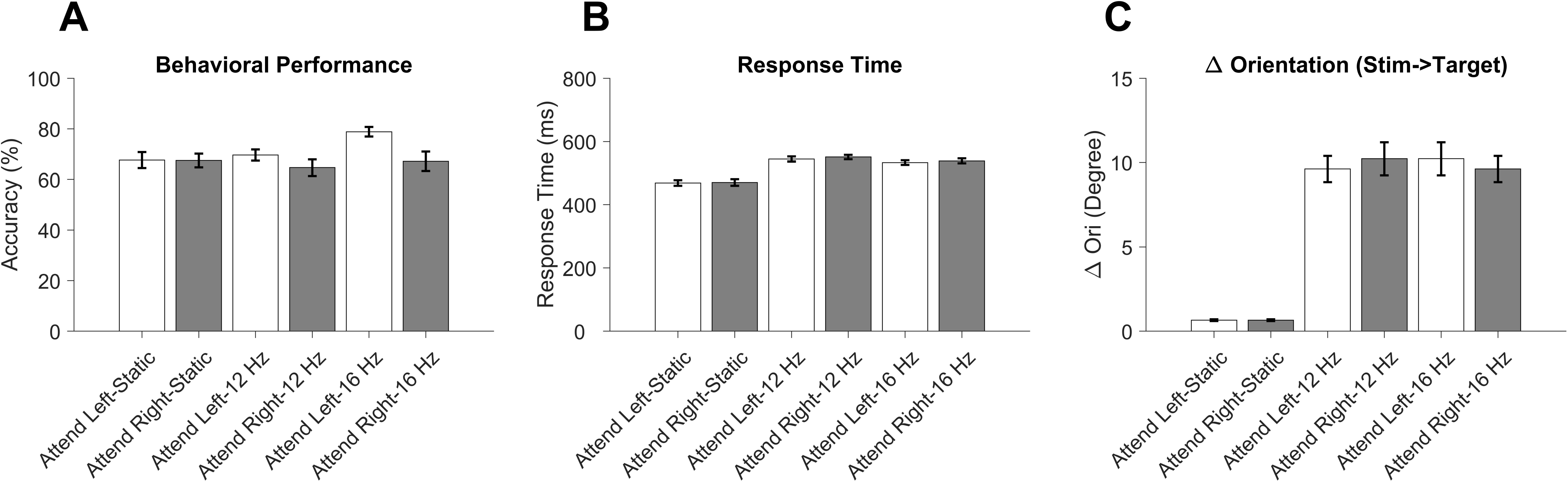
Behavioral Results for Attention Task. (A) Behavioral accuracy (in percent) for different attended locations (left-white bars, right-gray bars) and different temporal frequencies of the cued/attended stimuli. (B) Response time (in milliseconds) for different attention locations and temporal frequencies of the cued/attended stimuli. (C) Change in orientation values (target) for different attention locations and temporal frequencies of the cued/attended stimuli.

### Comparison of attentional modulation on alpha and gamma response for static stimuli

We first analyzed the pre-target alpha and gamma responses (−;1000 ms to 0 ms w.r.t the target onset) for static stimuli. We used the baseline (−;1000 ms to 0 ms w.r.t the stimulus onset in a trial) alpha power of the attend-right condition to subtract from both the attend-left and attend-right (which also means ignored-left) pre-target period. The scalp maps for change in alpha power w.r.t baseline are shown in Figure 5A for attend-left and attend-right conditions. The elliptical disks in front of each scalp map show the attended location (left or right) as filled, and the ignored location is shown as blank disks. Alpha power was suppressed in the analyzable electrodes contralateral to the attended location. For the attend-left condition, we found alpha suppression in the contralateral analysis electrodes on the right hemisphere and vice versa for the attend-right condition. However, the alpha suppression was not lateralized in attend-right conditions like attend-left conditions. However, when we computed the scalp map of the difference between attend-left and attend-right (i.e., ignored-left), we observed alpha suppression on the right-side analysis electrodes (contralateral to attended side) compared to the left electrodes (ipsilateral to the attended side), consistent with previous findings on the attention-mediated alpha suppression in the occipital regions in the contralateral hemisphere in both human EEG studies (Foxe et al., 1998; Sauseng et al., 2005; Rihs et al., 2007) and monkey LFP studies (Chalk et al., 2010; Das and Ray, 2018; Prakash et al., 2021).

**Figure 5:**
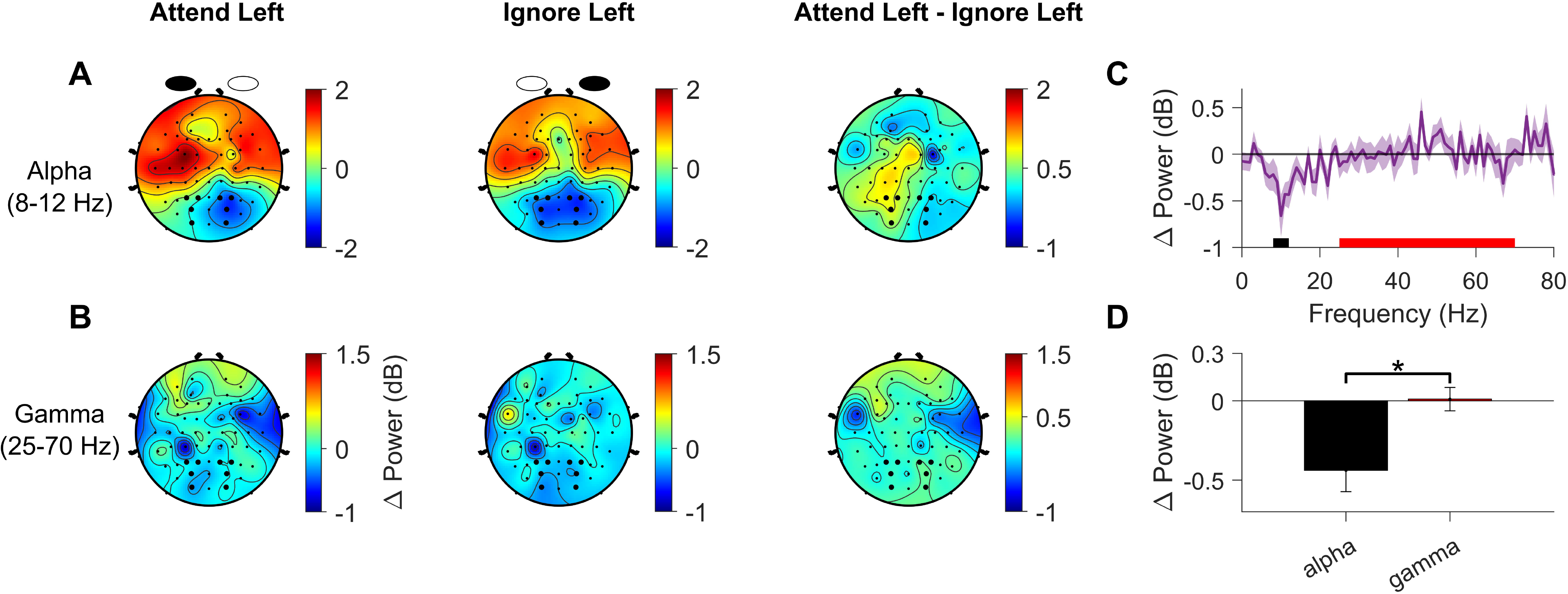
Attentional modulation of alpha and gamma response in attention tasks for static stimuli. (A) Scalp maps showing the change in power in the pre-target analysis period (−;1000 to 0 ms of target onset) w.r.t ignore left (i.e., attend-right) baseline (−;1000 to 0 ms of first stimulus onset in a trial) for alpha power (8-12 Hz) averaged across 26 subjects. The left column represents the attend-left condition, the middle column represents ignore-left (attend-right), and the right column indicates the difference between attend-left and attend-right conditions. Analysis electrodes used in Panels C and D are marked as bold dots. (B) Layout same as A for gamma power (25-70 Hz). (C) Change in PSD vs. frequency for the attended condition versus ignored condition in the pre-target analysis period (−;1000 to 0 ms of target onset averaged across all 26 subjects for the eight occipital analysis electrodes (difference between the 4 electrodes on the right and 4 electrodes on the left) highlighted in panel A. Solid black and red bars on the X-axis indicate alpha and gamma bands, respectively. (D) Change in alpha (black bar) and gamma (red) power between the attended condition and the ignored condition in the pre-target analysis period (−;1000 to 0 ms of target onset), averaged across all 26 subjects for the eight occipital analysis electrodes highlighted in panel A. The respective powers were calculated in the frequency ranges highlighted in panel B. For statistical comparison of the magnitude of attentional modulation between alpha and gamma power, we sign-reversed the alpha power values (negative values to positive and vice versa) and performed a pairwise t-test. Error bars indicate the standard error of the mean across subjects (N=26).

However, we found negligible effect of attention on gamma power in the scalp maps and the analysis electrodes (Figure 5B). We quantified the attention-modulated band power response for alpha and gamma for the occipital analysis electrodes (see method section). We found significant alpha suppression with attention (t-test, N= 26, p = 3.3 × 10^-3^) but negligible gamma modulation (t-test, N= 26, p = 0.88), with significant difference in attentional modulation between alpha and gamma magnitudes (Figure 5D; t-test, N= 26, p = 0.03).

### Comparison of attentional modulation of alpha, gamma, and SSVEP response for flickering stimuli

For the SSVEP response for the flickering condition, four conditions had to be combined since the attention task had two attended locations (left or right) and two different frequency tags (12 and 16 Hz). Figure 6A-D shows the scalp maps for all these four conditions. The filled and blank elliptical disks in each scalp map indicate the attended location (left or right), and the ignored location, respectively, with the flickering frequency tags mentioned on top of the disks. As expected, the SSVEP responses for all combinations were higher for the attended condition than the ignored condition for the contralateral occipital electrodes, consistent with an earlier study (Joon Kim et al., 2007; see their Figure 3). We consolidated all the SSVEP responses for a simple and effective comparison across different neural measures by presenting SSVEP responses on the right hemispheric occipital electrodes for a consolidated scalp map as if the attended location was always on the left. To achieve this, we mirror-flipped the scalp maps with respect to the midline set of electrodes (z-line of ActiCap-64 Cap) for the attend right conditions (Figure 6B and Figure 6D), and then averaged the four conditions (Figure 6E).

**Figure 6:**
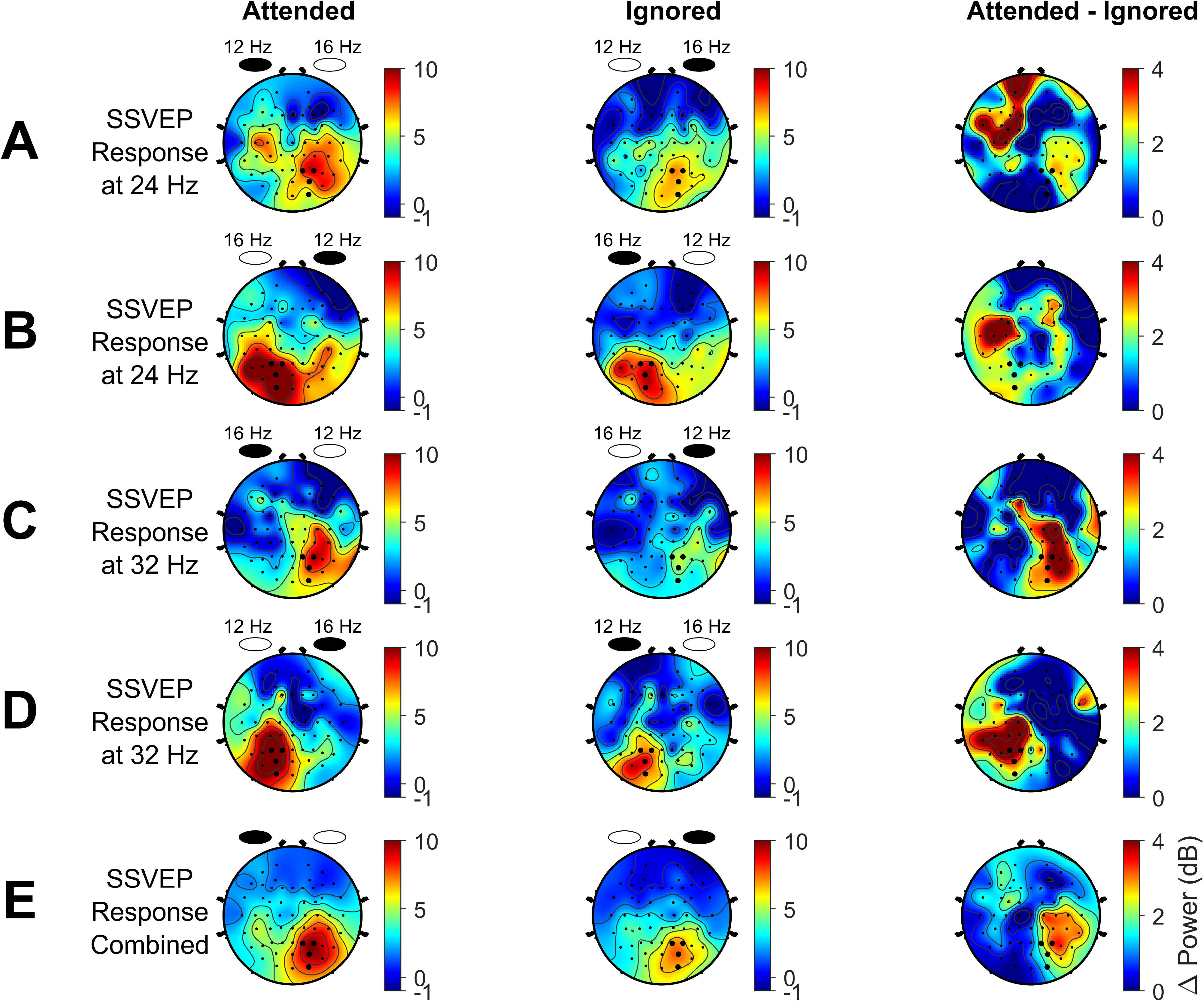
Scalp maps for attention-mediated SSVEP modulation for different attended locations and temporal frequencies of stimuli. (A-D) The attended location in each condition is presented as filled elliptical disks and the ignored location as blank elliptical disks. The flickering frequencies are mentioned on top of the disks. The analysis electrodes are highlighted in bold. (E) Consolidated scalp map after averaging responses in panels A-D. For panels B and D, the scalp maps were mirror-flipped w.r.t the midline of the cap to yield SSVEP responses in the right hemisphere prior to averaging.

Finally, we compared the attentional modulation of alpha, gamma, and SSVEP responses for flickering stimuli. We combined the alpha responses for both 12 Hz and 16 Hz flickering stimuli. Similar to the static stimuli, we found significant alpha suppression in the analyzable occipital electrodes in the contralateral hemisphere to the attended location (Figure 7A, top row; t-test, N= 26, p = 2.32 × 10^-6^). Interestingly, we found more substantial alpha suppression for flickering stimuli (mean ± SEM: −0.69 ± 0.11 dB across 26 subjects) compared to the static stimuli (−;0.43 ± 0.13 dB), potentially because the flickering stimuli produced stronger and more sustained activation of the cortex compared to the static stimuli. Again, attentional modulation of gamma was negligible for flickering stimuli, as can be observed in the scalp maps (Figure 7A, second row) and bar plots (Figure 7C, t-test, N= 26, p = 0.09). We computed SSVEPs using two methods (please refer to the Methods section) and combined different conditions (as discussed previously). SSVEP responses were enhanced in the trial-averaged estimate (Figure 7A, the bottom row) compared to the trial-by-trial estimate (Figure 7A, third row). This can be better appreciated by comparing the SSVEP power at 24 and 32 Hz in the trial-by-trial PSD estimates (Figure 7B, top) with the trial-averaged PSD (Figure 7B, bottom). We used data from 19 subjects for this panel, since for seven subjects, the SSVEP response appeared at 23 Hz and 31 Hz instead of the expected 24 and 32 Hz because of an extra stimulus frame during the presentation of flickering stimuli. Overall trends for these seven subjects were similar except the SSVEP peaks were at 23 and 31 Hz. For scalp map and bar plot analyses, we pooled the responses at 23 and 31 Hz for seven subjects and 24 and 32 Hz for the remaining 19 subjects. The SSVEP response at 24 and 32 Hz (cyan lines) improved when we computed the trial-averaged SSVEP, although the standard deviation was also higher.

**Figure 7:**
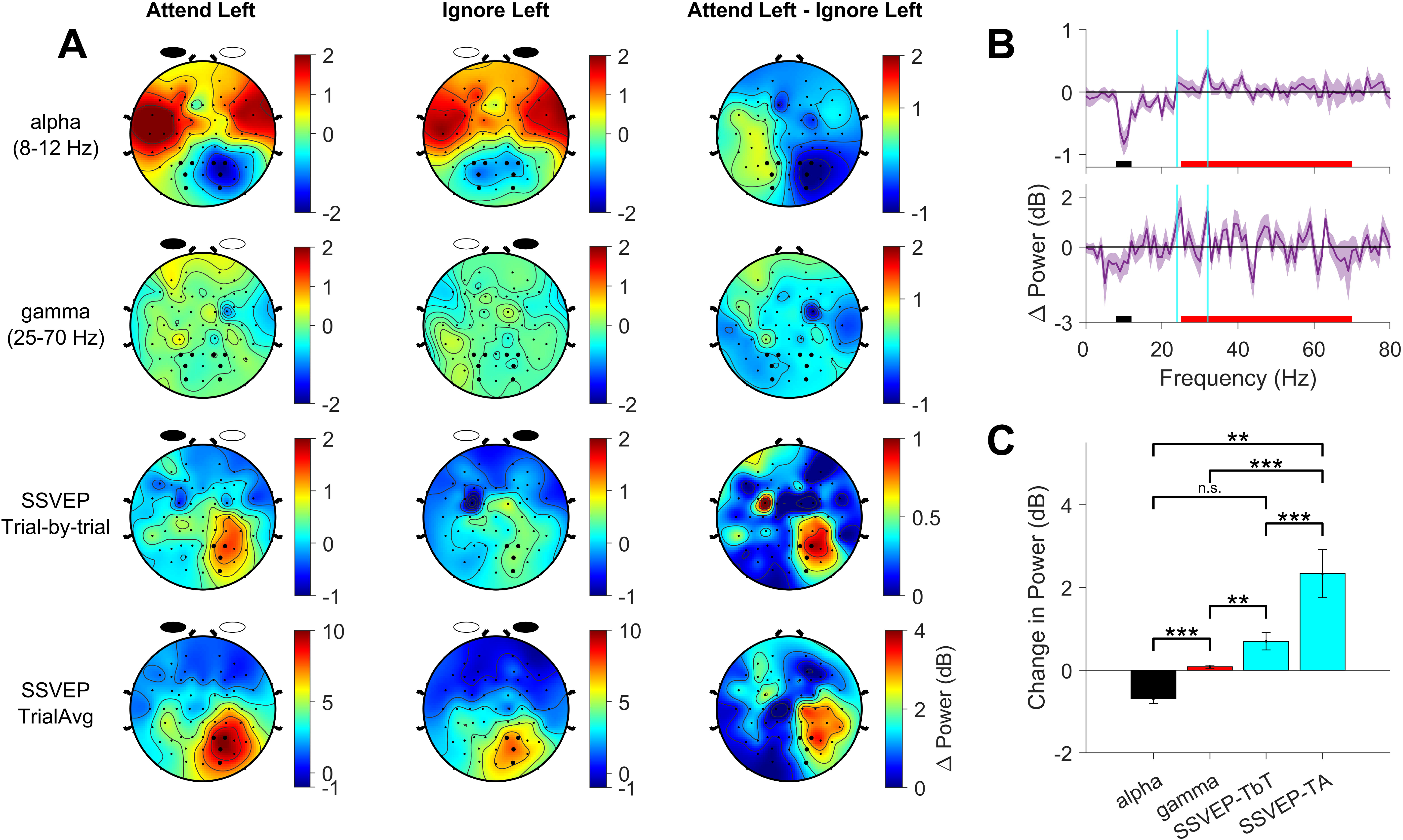
Modulation of alpha, gamma, and SSVEP response in attention tasks for flickering stimuli. (A) Scalp maps showing the change in alpha (8-12 Hz), gamma (25-70Hz), trial-by-trial SSVEP power, and trial-averaged SSVEP power (combined for 24 and 32 Hz) (top to bottom row) in the pre-target analysis period (−;1000 to 0 ms of target onset) w.r.t attend-right (i.e., ignored left) baseline (−;1000 to 0 ms of first stimulus onset in a trial), averaged across 26 subjects. The left column represents the attend-left condition, the middle column represents ignored-left, and the right column indicates the difference between attend-left and ignored-left conditions. Analysis electrodes are highlighted. As explained in Figure 3, alpha/gamma and SSVEP responses are computed using different approaches, which is reflected by showing only the contralateral 4 electrodes for the SSVEP responses. (B) (top) Trial-by-trial estimate of change in power spectral density (PSD) vs. frequency for the attended condition and ignored condition in the pre-target analysis period (−;1000 to 0 ms of target onset), averaged across 19 subjects for the eight occipital analysis electrodes; (bottom) shows the same for trial-averaged signal estimates. Solid black and red bars on the X-axis indicate alpha and gamma bands, respectively whereas the SSVEP responses at 24 Hz and 32 Hz are highlighted by solid cyan vertical lines. (C) Change in alpha (black bar), gamma (red), and SSVEPs for both estimates (cyan) power between the attended and ignored condition in the pre-target analysis period (−;1000 to 0 ms of target onset), averaged across all 26 subjects for the eight occipital analysis electrodes. The respective powers were calculated in the frequency ranges highlighted in panel B. To statistically compare the magnitude of attentional modulation between alpha power and other neural measures, we sign-reversed the alpha power values (negative values to positive and vice versa) and performed a pairwise t-test. Error bars indicate the standard error of the mean across subjects (N=26).

We found a strong modulation in SSVEP response with attention (t-test, N= 26, p = 2.43×10^-3^) which was enhanced even more (t-test, N= 26, p = 4.48×10^-4^) if we computed SSVEP using the ‘trial-averaged estimate.’ There was a significant difference between attention-mediated alpha and gamma modulation (Figure 7C t-test, N= 26, p =1.61×10^-4^; note that alpha values were sign reversed before statistical analysis) as well as for gamma and trial-by-trial SSVEP modulation (t-test, N= 26, p = 6×10^-3^). The modulation in alpha was comparable to the trial-by-trial SSVEP estimate (t-test, N= 26, p = 0.96) but significantly different from the trial-averaged SSVEP estimate (t-test, N= 26, p = 6.6×10^-4^). Taken together, SSVEP and alpha outperformed gamma in capturing attentional modulation in human EEG studies.

## Discussion

We used both static and flickering stimuli to record three relevant neural measures: alpha power, gamma power, and SSVEP power in human EEG experiments under similar behavioral conditions and compared the magnitude of attentional modulation of those measures. Non-foveal stimuli did not produce robust gamma response in spite of several stimulus optimizations. Alpha and SSVEP were comparable and outperformed gamma in capturing the attentional effect in human EEG. Finally, the effect of attention was more prominent for trial-averaged SSVEP compared to trial-by-trial SSVEP.

### Attention studies at different recording scales - macaque LFP vs. human EEG studies

#### Alpha and Gamma Power

The effect of attention on alpha and gamma power in LFP signals has been studied extensively in multiple visual areas along the visual pathway, such as in V1 (Chalk et al., 2010; Das and Ray, 2018) and V4 (Fries et al., 2001; Prakash et al., 2021), as well as in human EEG studies (Klimesch et al., 1998; Worden et al., 2000; Sauseng et al., 2005; Thut et al., 2006). A few studies found attentional modulation of stimulus-induced gamma band activity in human EEG by introducing a unique stimulus and task paradigm where subjects had to attend to a moving bar or other moving stimuli on the left or right visual field (Gruber et al., 1999; Müller et al., 2000), which were different from Posner-spatial cueing task paradigm. We however used non-foveal grating stimuli in our task paradigm to remain consistent with classical spatial attention studies in macaques. Previous macaque LFP and human EEG studies have reported that gamma rhythms are stimulus-induced and depend on various stimulus properties such as spatial frequency, orientation, size of the stimuli (Ray and Maunsell, 2010; Jia et al., 2013; Murty et al., 2018) as well as eccentricity of the stimulus in human EEG/MEG studies (Muthukumaraswamy and Singh, 2013; van Pelt and Fries, 2013). We designed a systematic approach to find the optimum size and eccentricity of grating stimuli that would produce robust gamma response without compromising the difficulty of our covert spatial attention task but failed to elicit strong gamma response or gamma modulation with attention. This could be due to three reasons. First, since we used peripheral gratings, we expected gamma response to be reduced due to reduced cortical magnification factor for peripheral stimuli (Daniel and Whitteridge, 1961; Slotnick et al., 2001). Indeed, we found negligible stimuli-induced gamma response in all four size-eccentricity mapping protocols studies (Figure 3), although we used peripheral gratings of similar size and eccentricities used by van Pelt and Fries in their MEG study (2013). Second, gamma is a “local” signal which is coherent only over a short spatial scale (see Mechanisms section below for more discussion) and therefore may not be well represented in the diffuse neural activity recorded in EEG. Third, the effect of attention on gamma power is variable across visual cortex, with reduction in power in V1 and enhancement in V4 (Chalk et al., 2010). These effects may cancel out in an aggregate signal such as EEG, leading to negligible overall attentional modulation.

#### SSVEP Power

A recent macaque LFP study (Prakash et al., 2021) in area V4 found that alpha and gamma outperformed SSVEP in capturing attentional modulation, possibly due to the SSVEP frequency (10 Hz counterphase) being close to the alpha range, as shown in previous studies (Gulbinaite et al., 2019). Attention studies using human EEG have specifically used SSVEP power to study attentional modulation (Morgan et al., 1996), to understand sensory gain in the context of attention (Kim et al., 2007; Itthipuripat et al., 2014a), and to test out predictions of the normalization model of attention on neural gain mechanisms (Itthipuripat et al., 2014b). We also found an increase in SSVEP power with attention which was higher if SSVEP was computed on the trial-averaged signal compared to trial-by-trial basis. While alpha suppression and SSVEP enhancement are routinely observed in EEG studies, to our knowledge, no prior study has systematically measured the relative modulation by attention under matching stimulus and behavioral conditions.

### Neural mechanisms of alpha and gamma

Our results, together with our previous report using LFP (Prakash et al., 2021) in which attentional modulation was strongest for gamma frequencies and beyond, provide important clues about the spatial scales over which mechanisms related to various oscillations operate. For example, previous studies have pointed out a thalamo-cortical origin of occipital alpha oscillations with conflicting evidence of alpha pacemakers being present either in the thalamus (Saalmann et al., 2012; Vijayan and Kopell, 2012) or the cortex (Bollimunta et al., 2008; van Kerkoerle et al., 2014). For example, macaque LFP studies show higher spike-field coherence in the alpha range in the deeper layers compared to the superficial layers during attention tasks with attentional suppression of alpha power and coherence being stronger in deeper layers compared to superficial layers (Buffalo et al., 2011), which points towards the layer 5 alpha pacemakers. Since EEG signals capture a spatiotemporally smoothed version of the LFP signals, integrated over an area of 10 cm^2^ (Buzsáki and Wang, 2012) or more which is several orders of magnitude higher than that of LFP, the alpha pacemakers in the visual cortex and thalamus must be coherent over a long range to yield a prominent occipital alpha in EEG. In addition, there is evidence of alpha acting as a short-range inter-areal feedback signal among visual areas, as found from ECoG recordings (Halgren et al., 2019), human MEG recordings (Michalareas et al., 2016) as well as laminar recordings from multiple macaque visual areas (van Kerkoerle et al., 2014). Top-down attention signals from higher-order areas can therefore modulate the gain of lower-order areas by influencing the alpha feedback signals. Studies of spatial attention have demonstrated that when covert attention is directed to one hemifield, alpha power in the contralateral hemisphere is suppressed in response to behaviorally relevant stimuli whereas alpha power increases in the ipsilateral hemisphere encoding for irrelevant stimuli as evidenced by human EEG studies (Worden et al., 2000; Thut et al., 2006; Rihs et al., 2007) and macaque LFP studies that show alpha suppression for attend-inside RF versus attend-outside RF conditions (Fries et al., 2001; Chalk et al., 2010). Together, these results suggest that the alpha rhythm is coherent over a relatively large cortical area, spanning perhaps a hemifield, such that robust alpha suppression can be observed even in an aggregate signal such as EEG.

On the other hand, gamma oscillations are generated due to excitatory-inhibitory interactions (Buzsáki and Wang, 2012), especially involving soma-targeting parvalbumin positive interneurons (Bartos et al., 2007; Cardin et al., 2009; Sohal et al., 2009), which remain coherent only over a few millimeters (Jia et al., 2011; Murty et al., 2018), implying that gamma oscillations are local in nature. Unlike alpha, gamma is shown to act as feedforward signals in visual cortex (van Kerkoerle et al., 2014; Michalareas et al., 2016) and gamma generated in local neural networks can interact with each other through synchronization to propel the signals from lower-order visual areas to higher-order visual areas. Cortical spread of LFP signals can capture gamma in the local population and the effect of attention can also be captured in the LFP signals possibly due to comparable spatial spread of LFP (∼0.5-1 mm; Katzner et al., 2009; Xing et al., 2009; Dubey and Ray, 2016) to that of attentional field (∼0.5-2.5 mm; Connor et al., 1997; Womelsdorf et al., 2008) in the visual cortex (see Prakash et al., 2022 for further discussion). The gamma generated in the local population in response to peripheral stimuli could have possibly been lost, as observed in our study (Figure 3), due to spatial averaging and reduced cortical magnification factor. However, robust gamma response has been reported in LFP gamma studies (Ray and Maunsell, 2010; Jia et al., 2013) or attention studies (Fries et al., 2001, 2008; Chalk et al., 2010) when non-foveal stimuli presented inside receptive field of microelectrodes. MEG signals, with better spatiotemporal resolution (∼2-3 mm; Buzsáki and Wang, 2012), can also capture the gamma generated by peripheral stimuli (van Pelt and Fries, 2013) and attentional modulation of gamma (Bauer et al., 2014; Magazzini and Singh, 2018). Taken together, this implies that mechanisms responsible for the generation of gamma oscillations are too local to be well captured in EEG.

### BMI applications for decoding focus of attention

Previous LFP studies in macaque visual areas have shown spikes and LFP can be reliably used to decode attentional state or attended location (De Sousa et al., 2021; Prakash et al., 2021). On the other hand, our study shows that the effect of attention was equally prominent in alpha and SSVEP power, making them more useful measures in decoding attention using human EEG. In fact, the effect of covert spatial attention on occipital alpha power is so salient in EEG across contralateral versus ipsilateral hemisphere, even for single trials, that it has earlier been used in brain-machine interfaces (van Gerven and Jensen, 2009; Bahramisharif et al., 2010). SSVEPs, on the other hand, have been used extensively in BMI applications to decode the focus of attention based on stimulus location or feature using human EEG (Min et al., 2016; Chinchani et al., 2022). Although the underlying neural mechanisms might be different; since alpha and SSVEP both show comparable attentional modulation in EEG, a combination of both the neural measures may be able to decode attended location better than a single measure. Therefore, our findings can have important implications for development of better BMIs to decode attentional state.

## Acknowledgments

This work was supported by Wellcome Trust/DBT India Alliance (Grant IA/S/18/2/504003; Senior Fellowship to S.R.), Tata Trusts, and Department of Biotechnology-Indian Institute of Science (DBT-IISc) Partnership Programme. A.D. is thankful for the financial support received from a research fellowship awarded by IISc, funded by the Ministry of Education, Government of India.

